# Discovery of the First Human Retro-Giant Virus: Description of its morphology, retroviral kinase and ability to induce tumours in mice

**DOI:** 10.1101/851063

**Authors:** Elena Angela Lusi, Federico Caicci

## Abstract

**Background:** The study of retroviruses dates back to the early 1900s during investigations on neoplastic diseases in chickens. Subsequently, Robert Gallo reported the first human retrovirus HLTV in 1980. What we report here is not an archetypal retrovirus, but the discovery of an oncogenic giant microbial agent with a mega-genome, where the transforming retroviral nature co-exists with multiple archaeal oncogenes.

**Methods:** After their isolation from human T cells Leukaemia, these organisms were examined at electron microscopy, tested for reverse transcriptase activity, fully sequenced, used for transformation tests on NIH-3T3 cells in vitro and tumours formation in mice. Same type of particles were also isolated from Canine Transmissible Venereal Tumour (CTVT), the oldest contagious cancer in nature.

**Results:** EM showed the presence of giant viral particles displaying retroviral antigens. These microbial entities harbour in their mega-genome a transforming retroviral kinase, cell-based oncogenes and have reverse transcriptase activity. The purified viral particles transformed NIH-3T3 cells and induced metastatic tumours in nude mice, three weeks post infection. Ruling out the possible presence of filterable retroviruses, a filtered supernatant did not display RT activity and did not transforms.

**Conclusions:** We discovered an ancestral microbial agent, acutely transforming. For its giant dimensions, the ability to retain the Gram stain, the presence of a mega-genome and its retroviral nature, we tentatively named the agent Retro-Giant-Virus (RGV). However, distinct from amoeba giant Mimiviruses, this transforming human agent has a different nature and does not require for its isolation amoeba co-culture, since amoeba is not its natural host.

The morphology, biology and genetic features allocate this mammalian giant microbe halfway in between a classic oncogenic virus and an infectious cancer cell. Its transforming nature goes with its constant ability to induce tumours formation in mice.

## Introduction

Our previous paper described the presence of unusual Mimiviruses-like structures in human tissues ^1^. Like Mimiviruses (∼450 nm giant viruses found in the amoeba), these human structures had the ability to retain Gram staining, and mass spectrometry revealed the presence of histone peptides that had the same footprints as giant viruses ^2–8^. However, the human giant virus-like structures displayed a distinct and unique mammalian retroviral antigenicity.

Our initial discovery in human tissues presented the conundrum of whether the structures were giant viruses with a retroviral nature or cellular components having a viral footprint. The distinction between the virus and the cells was blurred. The most difficult part to explain arose from the unique mammalian retroviral antigenicity associated with the human Mimiviruses-like structures. The gigantic dimension of our particle excluded it from the orthodox understanding of retroviruses but simultaneously presented the antipodean challenge to establish if this giant virus’s retroviral properties signified the discovery of the first human Retro-Giant Virus (RGV).

There was only one possibility to solve the dilemma: isolate the viruses (if really present) and verify if they contain genetic material. Consequently, we chose the traditional way of isolating virus using a sucrose gradient, but this time we did not use the common filtration techniques since they cause the loss of giant microbial agents.

In fact, classical virus isolation procedures include filtration through 0.2-µm-pore filters to remove bacteria and protists. However, these filters also often exclude large microbes. The recent discovery of Mimiviruses infecting amoeba highlighted how ad example the use of a 0.45 μm filter may prevent the recovery of giant viruses ^9^. Therefore, we adopted some of the strategies currently in use for the isolation of amoeba giant viruses, but the harvest of human giant microbes from cancer specimens did not require amoeba co-cultures.

In this manuscript, we describe all the steps that led to the isolation from human T cell leukaemia of a microbial organism that appears to be halfway in between a classic oncogenic virus and an infectious cancer cell.

The morphology and genetic structure of the giant agent, its ability to induce tumours formation in mice and the strong similarity between the particles isolated from Canine Transmissible Venereal Tumour (CTVT) with the human ones, establish the existence of an ancestral self-governing microbial cell-like entity acutely transforming, infecting humans and other species.

## Methods

### Isolation of giant viral particles from HPB-ALL, MOLT-4 and Loucy T cells leukemia on sucrose gradient

2 × 10^8^ human (HPB-ALL, DSMZ, Germany), MOLT4 and Loucy (ATCC, USA crl 2629, lot. 63734317) T cell leukaemia grown at 37°C in RPMI-1640, 10% fetal bovine serum and 2 mM L-glutamine, were centrifuged at 1,500 rpm g for 5 minutes at 4°C. The cell pellet was washed with 1x PBS. The cells were lysed (vortexed) with 2.5 ml PBS in the presence of 25μl of protease inhibitor cocktail (Abmgood, Richmond BC, Canada). Cell suspension was vortexed and incubated a 4°C for 30 minutes. Cell lysis was monitored using a phase contrast light microscope. The resulting crude extract was centrifuged at 3,000 rpm for 5 minutes. The pellet containing the cellular nuclei was discharged. The resulting supernatant was collected and slowly dripped over 9 ml of a 35-30-25% sucrose gradient (Sigma, Milan, Italy) and centrifuged at 10,000 rpm for 2 h in a 15 ml Polyclear Centrifuge tubes (Seton USA). Once a visible white disk, corresponding to 25% sucrose fraction, was observed, the fraction was collected after centrifugation at 14,000 rpm for 30 min, at 4°C.

### Electron microscopy (EM) immunogold of the giant viral particles

25 μl of the 25% sucrose isolated fraction was placed on Holey Carbon film on Nickel 400 mesh. The grids were treated for 30 minutes at room temperature with the primary monoclonal antibody (moAb) anti-Feline Leukaemia Virus p27-gag (catalog number, PF12J-10A; Custom Monoclonals International, West Sacramento, CA, USA) and subsequently with a secondary anti-mouse gold conjugated antibody (BB international anti-mouse IgG 15 nm gold conjugate; catalogue number, EM.GMHL15, Batch 4838). After staining with 1% uranyl acetate, the sample was observed with a Tecnai G2 (FEI) (Thermo Fisher) transmission electron microscope, operating at 100 kV. Images were captured with a Veleta (Olympus Soft Imaging System) digital camera.

### Gram positive stain of the giant viral particles

Gram positive staining of purified human giant microbial agent was performed with Colour Gram 2 Biomerieux kit, following the manufactures instructions. Before staining, slides were heated fixed 3–4 times through the Bunsen burner flame.

### RNA extraction from the giant viral particles

The microbial pellet was lysed with 1ml of RNA-XPress Reagent (Himedia, Mumbai, India), a monophasic solution of phenol-guanidine thiocyanate, and incubated at room temperature (RT) for 5 minutes. This was followed by the addition of 200 µL chloroform, vortexing for 15 sec and incubation at RT for 10 min. The organic and aqueous phases were separated by centrifuging the sample at 11,000 rpm for 15 minutes at 4°C. The aqueous phase, containing RNA, was harvested and precipitated with 600 μl of isopropyl alcohol and glycogen. After incubation for 1h at 20°C, RNA was pelleted by centrifugation at 11,000 rpm for 10 minutes. The RNA pellet was washed with 75% of ethanol, air dried and resuspended in RNase free H_2_O (Himedia). One aliquot was utilized for concentration determination in a MaestroNano Spectrophotometer (Maestrogen Inc, Hsinchu City, Taiwan).

### Reverse transcriptase (RT) activity of the giant viral particles

After sucrose gradient isolation, the microbial pellet was lysed in 20 μl of 20 mM Tris-HCL pH7.5, 100 mM NaCl, 0.1 mM EDTA, 1mM DTT, 50% (v/v) glicerol, 0.25% Triton X-100 (Sigma). To test the ability of the human giant viruses to retro-transcribe, 10 μl of the viral lysate, instead of a reverse transcriptase enzyme, were used to retro-transcribe 1 μg of total RNA from Human Liver Total RNA (ThermoFisher Scientific, Waltham, MA, USA). The reverse transcriptase reaction for the viral pellet was carried out with random primers using a commercial kit (EasyScript cDNA Synthesis kit; Abmgood), deprived of the supplied reverse transcriptase enzyme. The reverse transcriptase reaction was carried at 25°C for 10 minutes, then at 42°C for 50 minutes. The reaction was stopped by heating at 85°C for 5 minutes. The viral reverse transcriptase activity was compared to positive controls where a commercial RT enzyme was included (EasyScript RTase; Amgood).

After the reverse transcription, 2 μl of the obtained single stranded cDNA was further amplified in presence of 10 pmol of primers for GAPDH, 1.25 units of thermostable DNA polymerase (Precision DNA Polymerase; Abmgood), 0.2 mM dNTPs/ 2.0 mM MgCl2 in 1X PCR buffer in a final volume of 25μl. PCR conditions were: 1 cycle of 95°C for 5 minutes; 40 cycles of 94°C for 1 minute, 58°C for 1 minute, 72°C for 1 minute; 1 cycle of 72°C for 5 minutes. 20 μl of the PCR reaction was loaded on a 1% agarose gel for electrophoresis.

### c-DNA synthesis

1 μg of total RNA was utilized for cDNA synthesis using EasyScript cDNA Synthesis Kit (Abmgood) according to the manufacturer’s instructions. Briefly, 20 μl of reaction contained 200 units of reverse transcriptase, 0.5 μM of random primers, 20 units of ribonuclease inhibitors, 500 μM dNTP. The reaction was carried out at 25°C for 10 min, then at 42°C for 50 min.

### Whole Genome Shotgun Sequencing

Sequencing and bioinformatics analysis were performed by Genomix4Life (University of Salerno). In details the libraries were prepared using DNA as starting material, with Nextera flex Kit (Illumina Inc). Libraries were sequenced (paired-end, 2 × 75 cycles) on NextSeq platform (Illumina Inc.) Fastq underwent to Quality Control using FastQC tool (https://www.bioinformatics.babraham.ac.uk/projects/fastqc/). *De novo* assembly was performed using Geneious (version 11.1) software. The assembler was used with standard parameters. Blast2Go was using to perform the blast alignment sequences www.blast2go.org. Annotation respect to NCBI Viruses database (taxa 10239) was made through Gene Ontology http://www.geneontology.org. The algorithm used was blastx-fast and the statistical significance threshold for reporting matches was set 1.0E-3. The number of sequence alignments to retrieve was set to 20. The other parameters was set as default. All statistical values regarding assembled contigs were computed using QUAST (QUAlity Assessment Tool).

### Focus Formation-NIH 3T3 cells Transformation Assays, post viral infection

NIH 3T3 cells (ATCC -lot 700009353) were propagated in DMEM plus 10% FBS. 3× 10^5^ NIH 3T3 cells per 10 cm culture dish were seeded with 10 ml of NIH 3T3 medium and incubated O/N at 37 °C, 5% CO_2_ for infection the next day. For the seeding cells were kept at 50-60% confluence. The following day, the medium, from NIH 3T3 cells to be infected, was aspirated and replaced with 9 ml of fresh regular NIH 3T3 medium, 1 ml of purified viruses aliquots and polybrene at a final concentration of 6 μg/ml. The infected cells were incubated O/N at 37°C, 5% CO_2_. Only one round of infection was performed and the virus-containing medium was replaced with regular NIH-3T3 medium. Uninfected NIH-3T3 cells and NIH 3T3 cells infected with a filtered supernatant were used as controls. Phenotypic changes were valuated every day, for 2-3 weeks and medium was replaced as necessary.

### Tumors formation in nude mice

For the *in vivo* tumorigenicity assay, six-to eight-week-old specific pathogen-free Hsd: Athymic Nude-Foxn1nu mice were provided by (Envigo RMS Srl, Zona Industriale Azzida 57, 33049 San Pietro Al Natisone, Udine (Italy) and maintained in an isolated clean room held at a regulated temperature (25 ± 2°C) and humidity (approximately 45–65%). The mice were housed under a 12 h/12 h light/dark cycle and fed ad libitum with rodent diet and water. All protocols were performed according to the Standard Operating Procedures and Animal Care Legislative decree N. 26/2014. Experimental proof of concept was conducted on five nude mice. Infected, neoplastic NIH 3T3 cells were inoculated intra-peritoneal (IP) in two mice and subcutaneously (SC) in one. In addition, pure giant particles microbial aliquots were directly injected in two mice, intra-peritoneal in one and intravenously in the other. Gross anatomy and pathological detection were performed at 3 weeks, 4 weeks and 5 weeks post infection. The animals were monitored daily for general clinical health conditions. Mice were euthanized by cervical dislocation, tissue samples were excised during necropsy and stored for downstream applications in formalin buffer 10%.

## Results

### Electron Microscopy of giant microbial particles isolated from human T leukemia cells

Giant particles isolated from human T cells leukemia (MOLT-4, HPB-ALL, Loucy) formed a white band at 25-30% interface of the sucrose gradient (that is also the sedimentation rate of environmental giant viruses infecting amoeba).

The band was recovered and examined at EM. An in depth EM morphological analysis revealed the presence of predominant ∼400-450 nm viral particles, that surround a much larger particle. These human *Giants* appear as an ancestral microbial system composed of ∼1-1.5μm large particle and smaller satellite associated viral particles, Mimiviruses-sized. Striking images of the giant microbes isolated from human T-cell leukemia are reports in Table 1, Table 2 and Figure 1. Same type of particles were also found in CTVT (Table 3). A comparative analysis of the morphological features between the human and CTVT giant particles suggests that this ancestral oncogenic microbial system is conserved and able to infect humans and other species, (Table 4).

**Table 1.**
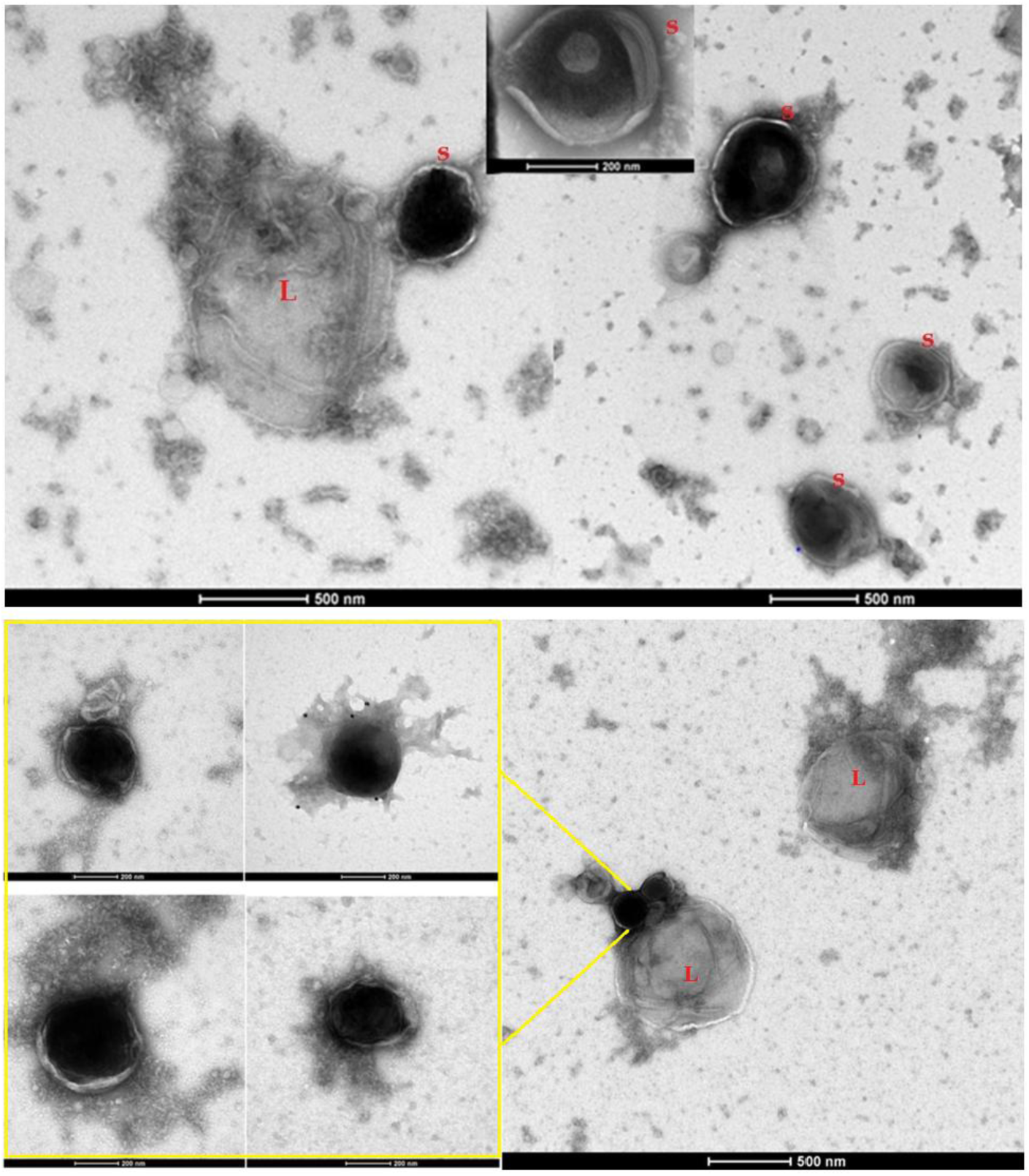
EM micrograph of giant viral particles purified from MOLT-4, HPB-ALL and Loucy human T cell leukemia, through a sucrose gradient. The predominant 400-450-nm satellite particles (S) surround a larger particle (L) of ≥1 micron. These biological entities appear as a microbial system composed by a unicellular organism and satellite smaller viral particles, Mimiviruses-sized.

**Table 2.**
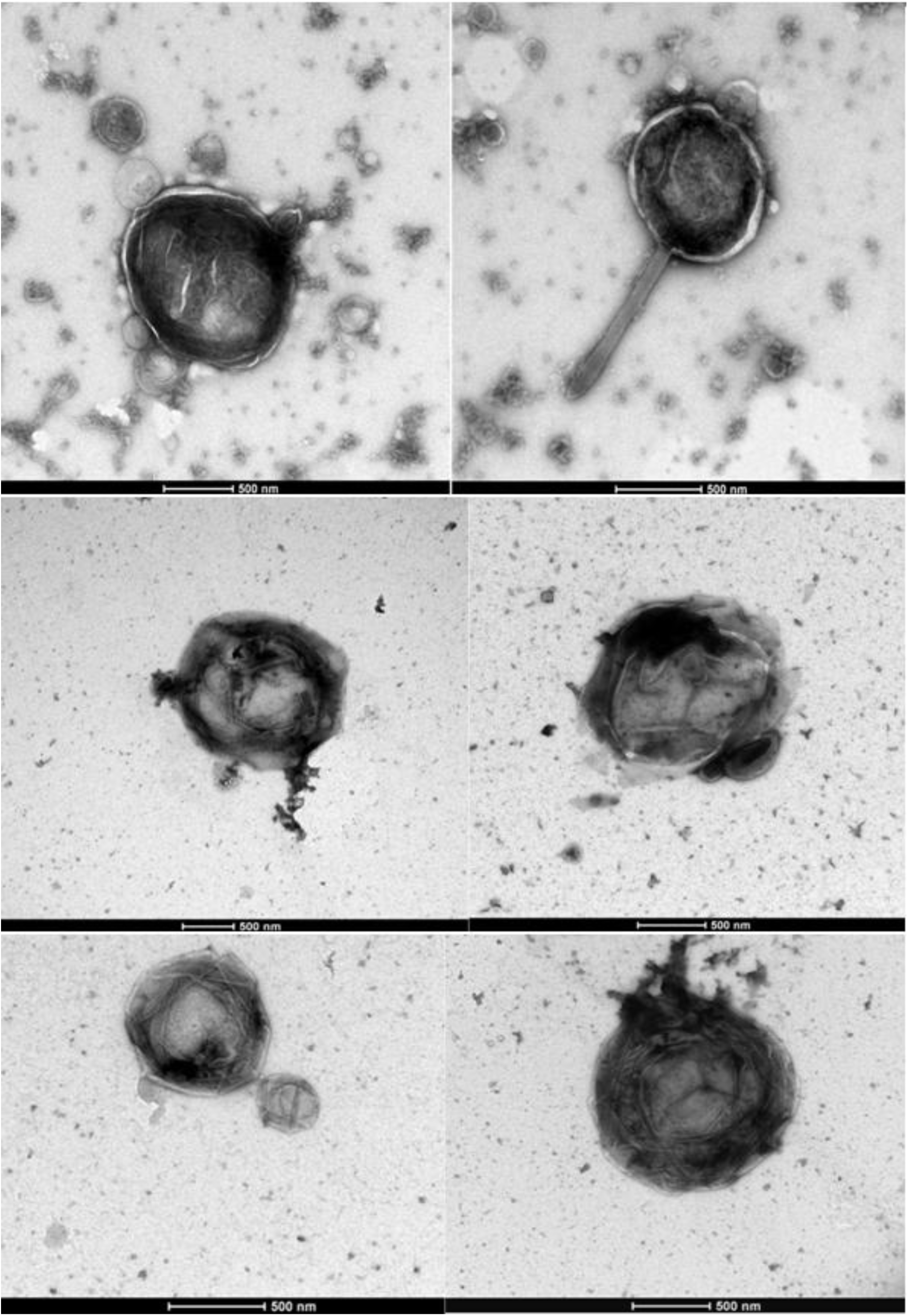
EM OF ≥ 1μm LARGE PARTICLE, isolated from human T cell leukemia, through a sucrose gradient. The morphology of the large particle that can reach up to 2 micron.

**Table 3.**
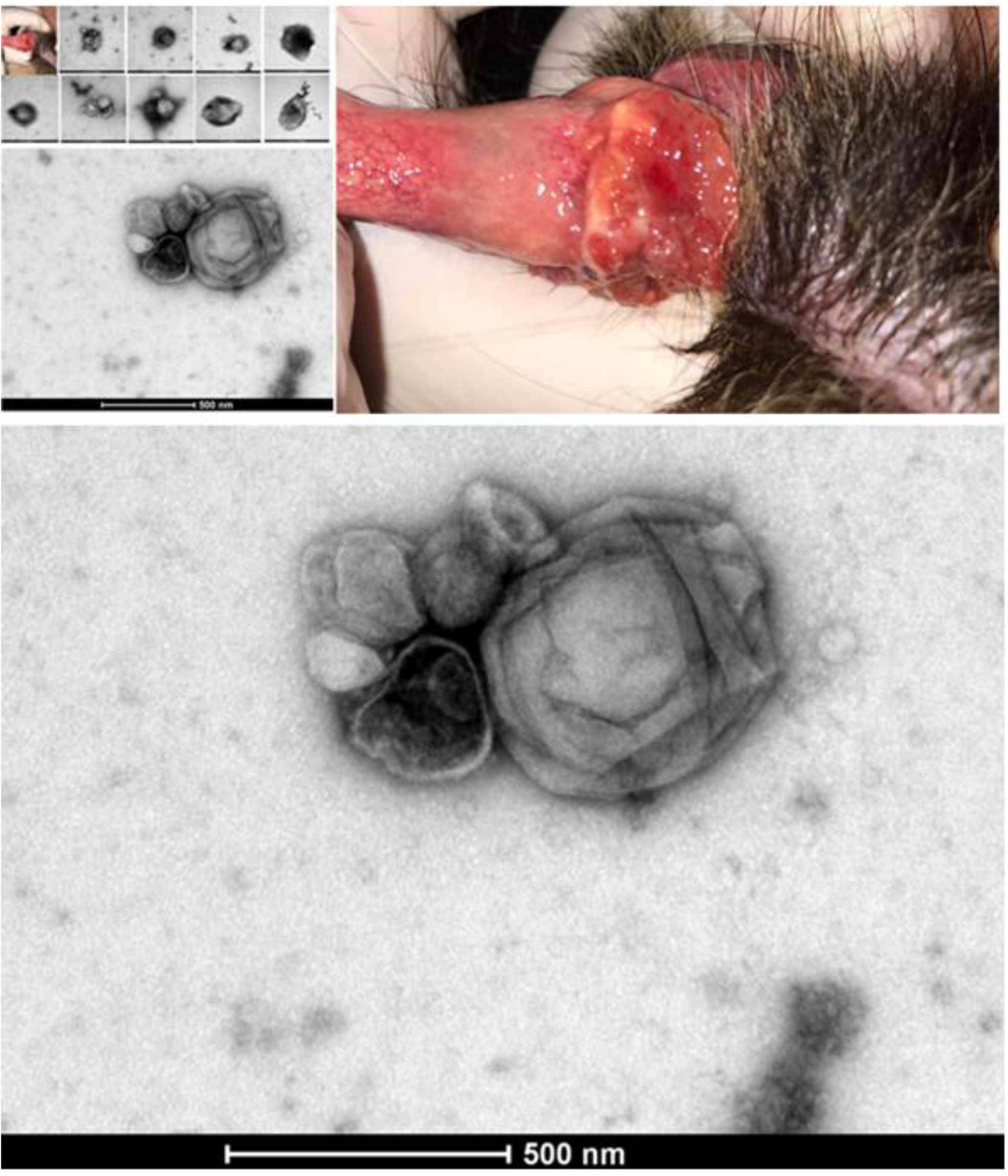
EM after sucrose gradient purification. Microbial giant particles isolated from CTVT. Same scenario as giant viral particles isolated from human leukemia cells. Smaller satellite 400-450 nm giant viral particles surround a large particle resembling ancestral unicellular organisms.

**Table 4.**
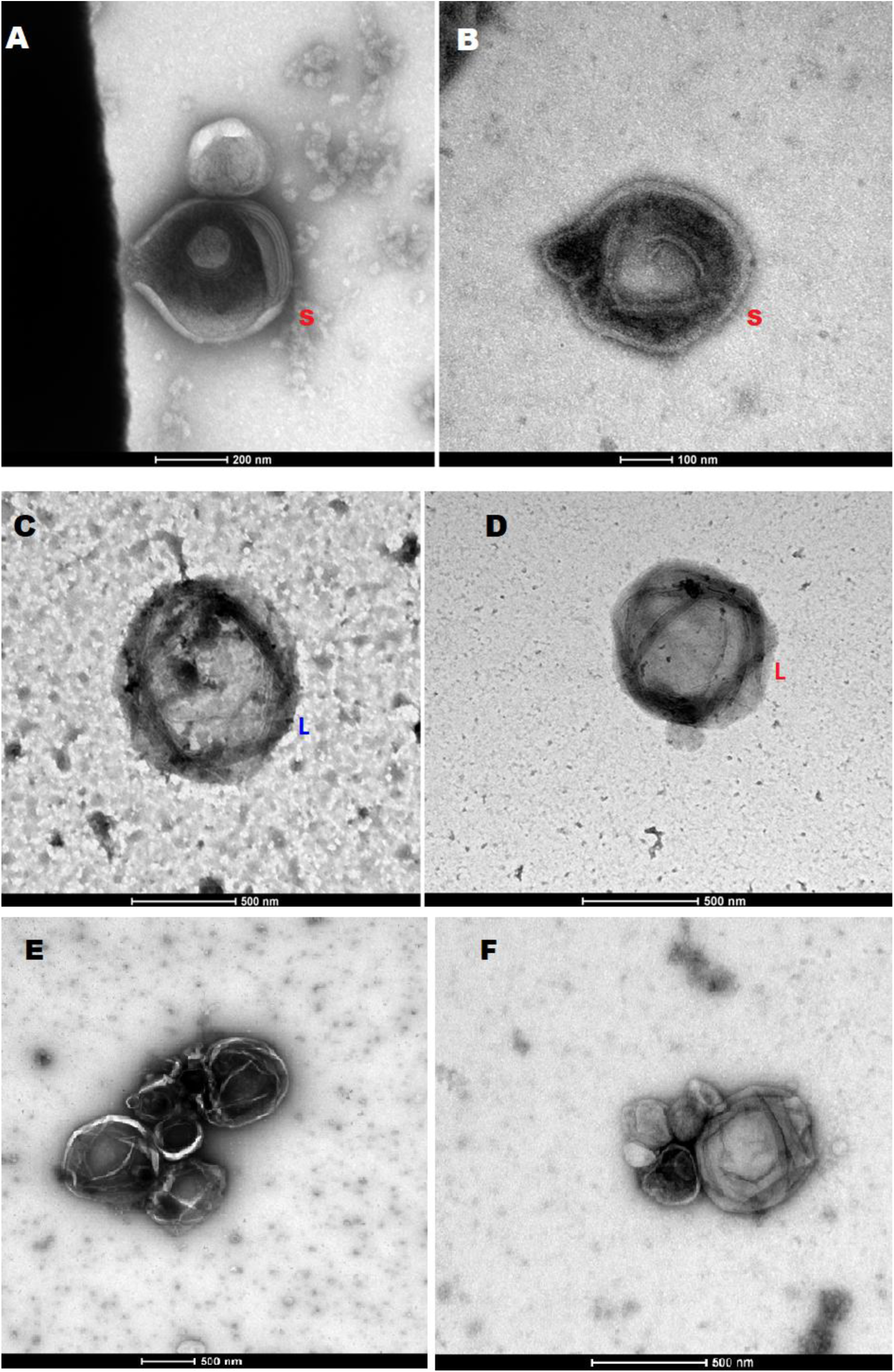
Comparative morphology between the human and CTVT giant viral particles. The large unicellular organism (L) and the smaller satellite giant viral particles (S) isolated from human neoplasms (human T cell leukemia) have the same morphology as the giant viral particles isolated from CTVT. Same scenario and same structure.

**Figure 1.**
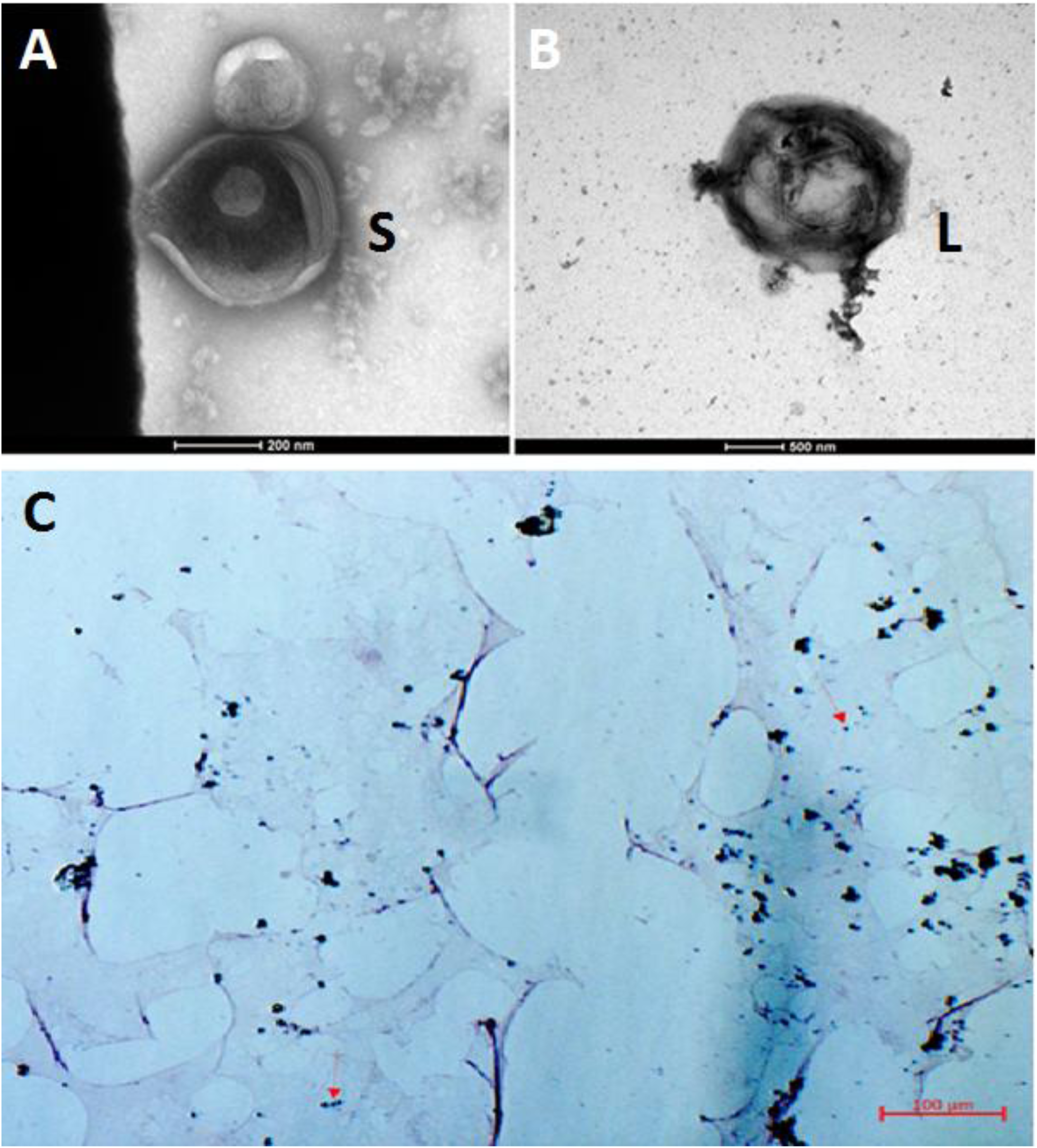
A) High resolution micrograph of the smaller (S) satellite viral particle, Mimiviruses sized. B) High resolution picture of the larger (L) particle that reaches in this micrograph 2 μm of dimension. C) Gram stain of the giant particles, after sucrose gradient purification. Like Mimiviruses infecting amoeba, these oncogenic human giants microbes have the ability to retain the Gram stain, but differently from Mimiviruses, their harvest do not require amoeba co-culture. Blue granules diagnose giant viruses (red arrows indicate some of these, but blue granules can be seen all over the slide).

### Focus Formation Assay-Transformation of NIH 3T3 Cells

These giant microbial entities behave as acutely rapid transforming viruses, showing both transforming and immortalizing effects. In NIH 3T3 cells, the loss of contact inhibition and initial foci formation start after 48 hours post infection.

In about a week, there were neoplastic foci in culture growing independently. The dynamics of malignant transformation in NIH 3T3 cells induced by these large agents is reported in Table 5 and Table 6.

**Table 5.**
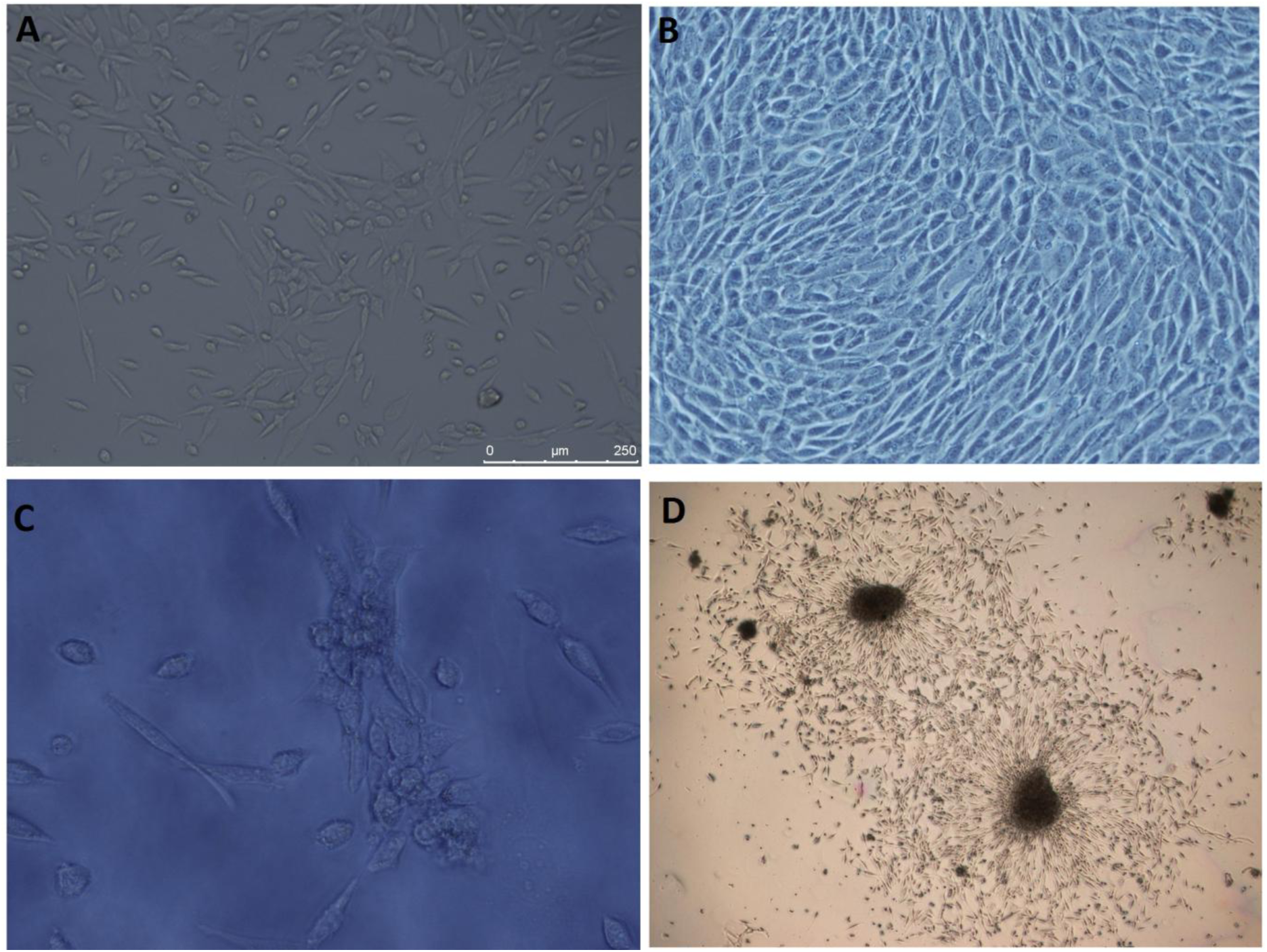
Focus formation assay on NIH 3T3 cells infected with purified aliquots containing the above described giant particles. A) Uninfected NIH 3T3 cells in culture with normal phenotype. B) NIH 3T3 cells infected with a filtered supernatant from human leukaemia cells with normal phenotype. C) NIH 3T3 cells 48 hours post giant particles infection, displaying loss of contact inhibition and initial foci formation. D) NIH 3T3 cells at day 10 post infection, with neoplastic foci growing independently in culture.

**Table 6.**
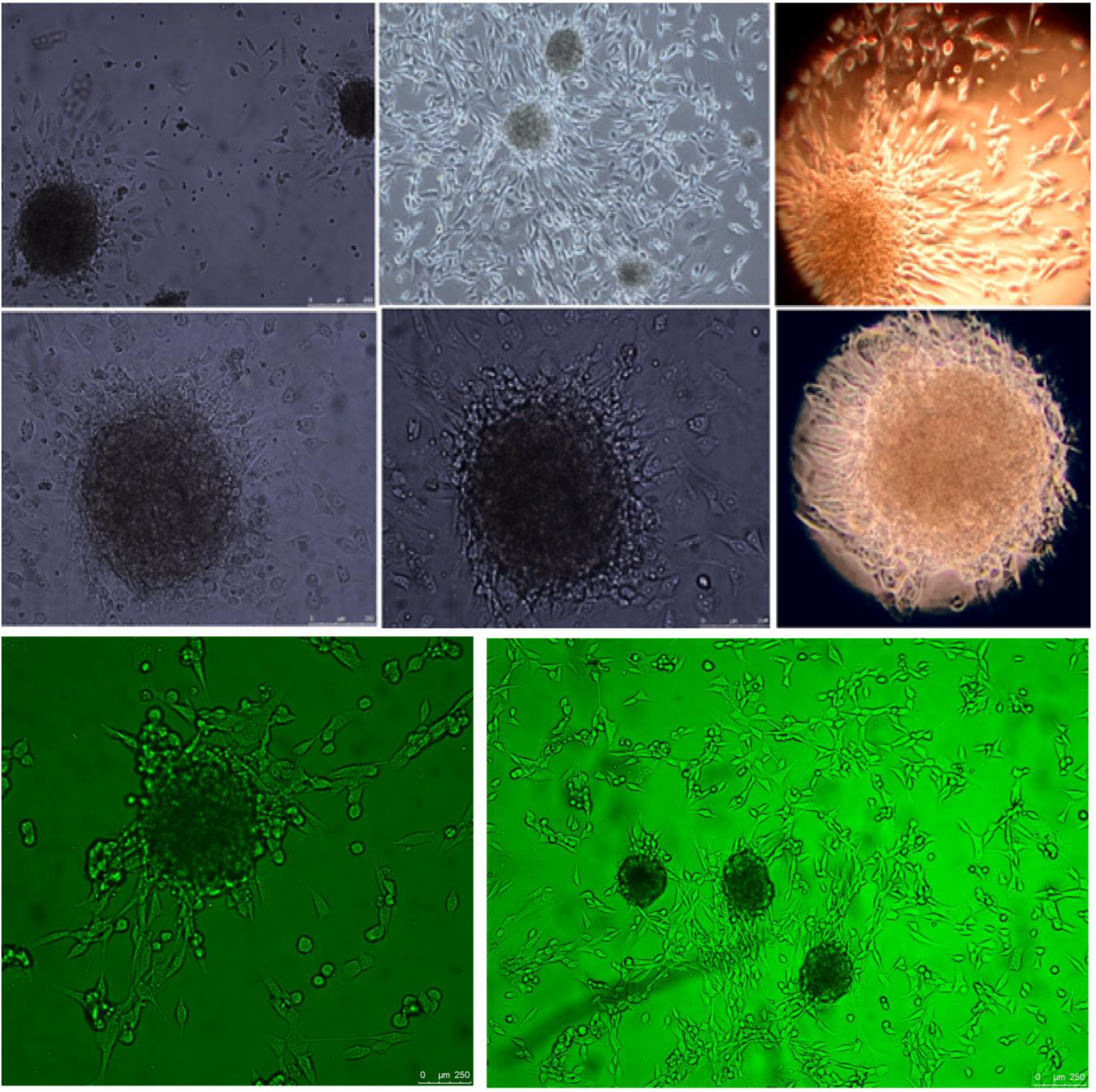
NIH 3T3 cells infected with human giant viral particles. Images of typical neoplastic FOCI at day ten post infection.

It should be noted that only the fraction containing the *Giants* induced neoplastic foci formation in NIH 3T3 cells, while a filtered supernatant did not. This ruled out possible role of filterable viruses or retroviruses.

Full filling the Koch’s postulates, the giants viral particle could be re-isolated from infected NIH 3T3 cells that became cancerous, with the usual routinely sucrose gradient.

### Tumorigenicity in Nude Mice

Experimental proof of concept was conducted on five nude mice. The characteristics of the animals, ethics and study design are detailed in the Materials and Methods section. Infected, neoplastic NIH 3T3 cells were inoculated intra-peritoneal (IP) in two mice and subcutaneously (SC) in one. In addition, pure giant particles microbial aliquots were directly injected in two mice, intra-peritoneal in one and intravenously in the other. Gross anatomy and pathological detection were performed at 3 weeks, 4 weeks and 5 weeks post infection. Four out of five mice developed macroscopically visible cancerous lesions within 3-5 weeks, (Figure 2). The histological diagnosis of the tumour explants confirmed the presence of poorly differentiated, highly invasive anaplastic fibro-sarcomas, Figure 3.

**Figure 2.**
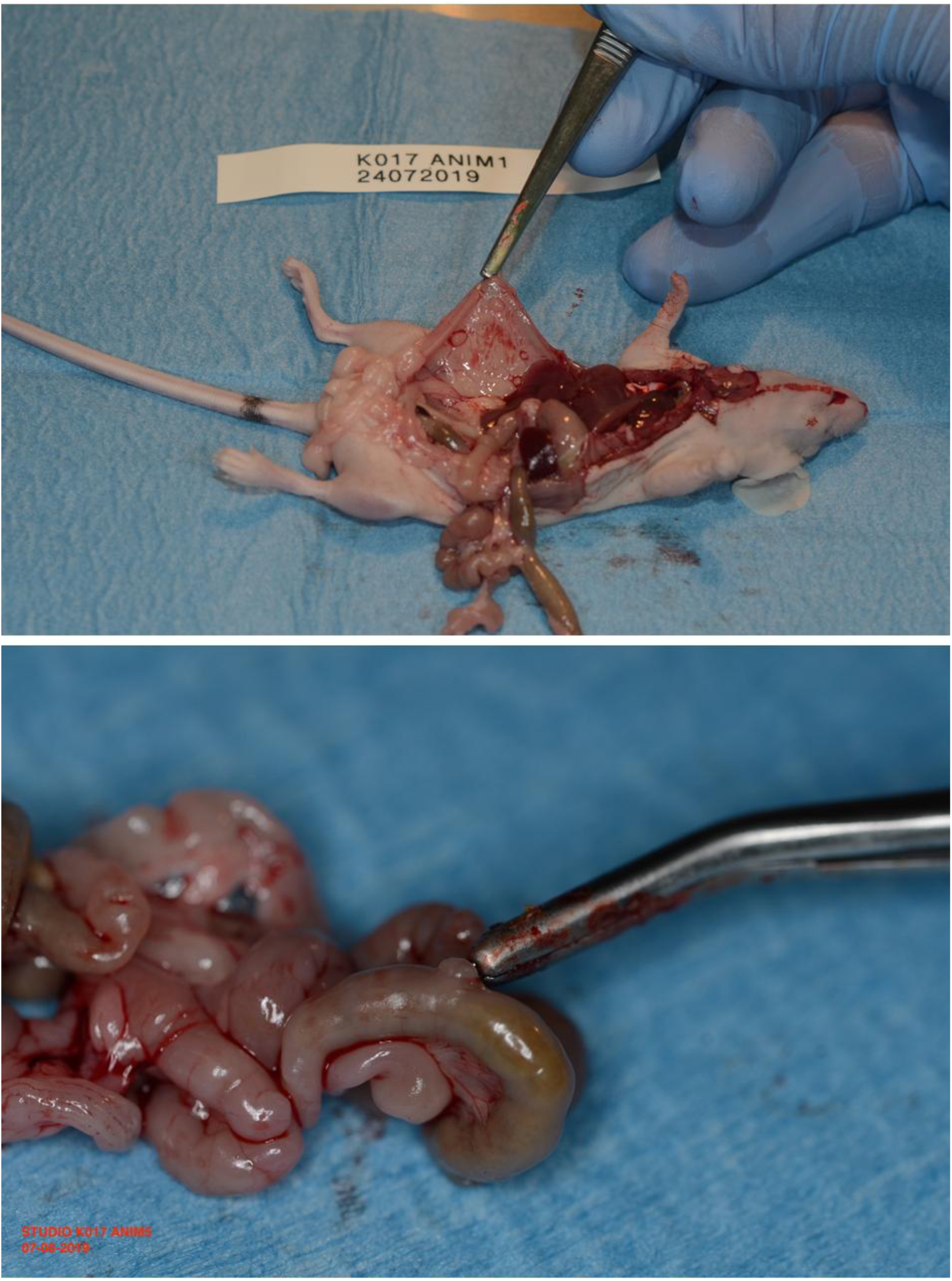
A) Multiple Peritoneal Nodules under the tweezers holding the omentum, three weeks post infection, route of infection IP. B) Scissor indicates an intestinal cancerous lesion, 5 weeks after infection with purified viral aliquot, route of infection IV.

**Figure 3.**
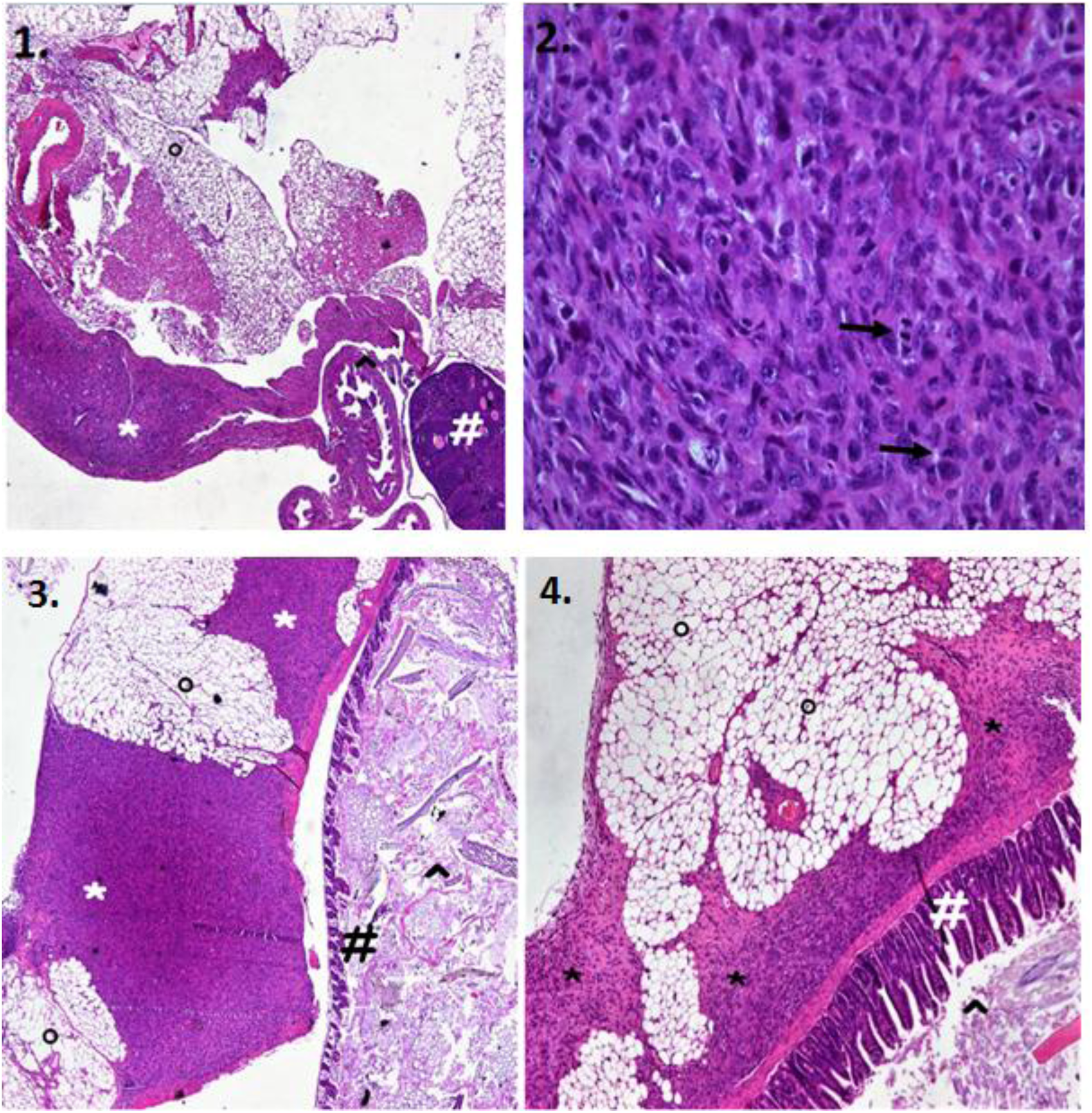
Section 1) °Peritoneal Adipose Tissue, * CANCER, ^ Fallopian tubes, # Ovary. Section 2) highly aggressive Anaplastic Fibro-sarcoma, magnification value X40. Black Arrows indicate mitotic figures. Section 3 and 4) # Intestinal Villi, ^ Intestinal lumen, °Peritoneal adipose tissue, * CANCER.

### Reverse transcriptase activity

These oncogenic giant particles have reverse transcriptase activity. 10 μl of the lysed purified viral pellet produced cDNA from an RNA template (Figure 4). It should be noted that HPB-ALL, MOLT-4 and Loucy leukaemia T cell lines, from which these giant agents were extracted, were screened by PCR and found EBV, HBV, HCV, HHV-8, HIV-1, HIV-2, HTLV-I/II, MLV, SMRV Negative. This ruled out co-infection with typical retroviruses. RT activity matched to the presence of retroviral antigens at EM immunogold. To test the presence of possible retroviral antigens expressed on the giant particles, we used anti-FeLV gag p27 moAb, used for its ability to bind conserved epitopes among different mammalian retroviruses ^10-16^. At EM immunogold, both the large and satellite ∼400 -450 nm giant viral particles shared the same retroviral nature, being specifically labelled by anti-FeLV gag p27 mo-Ab, (Figure 5).

**Figure 4.**
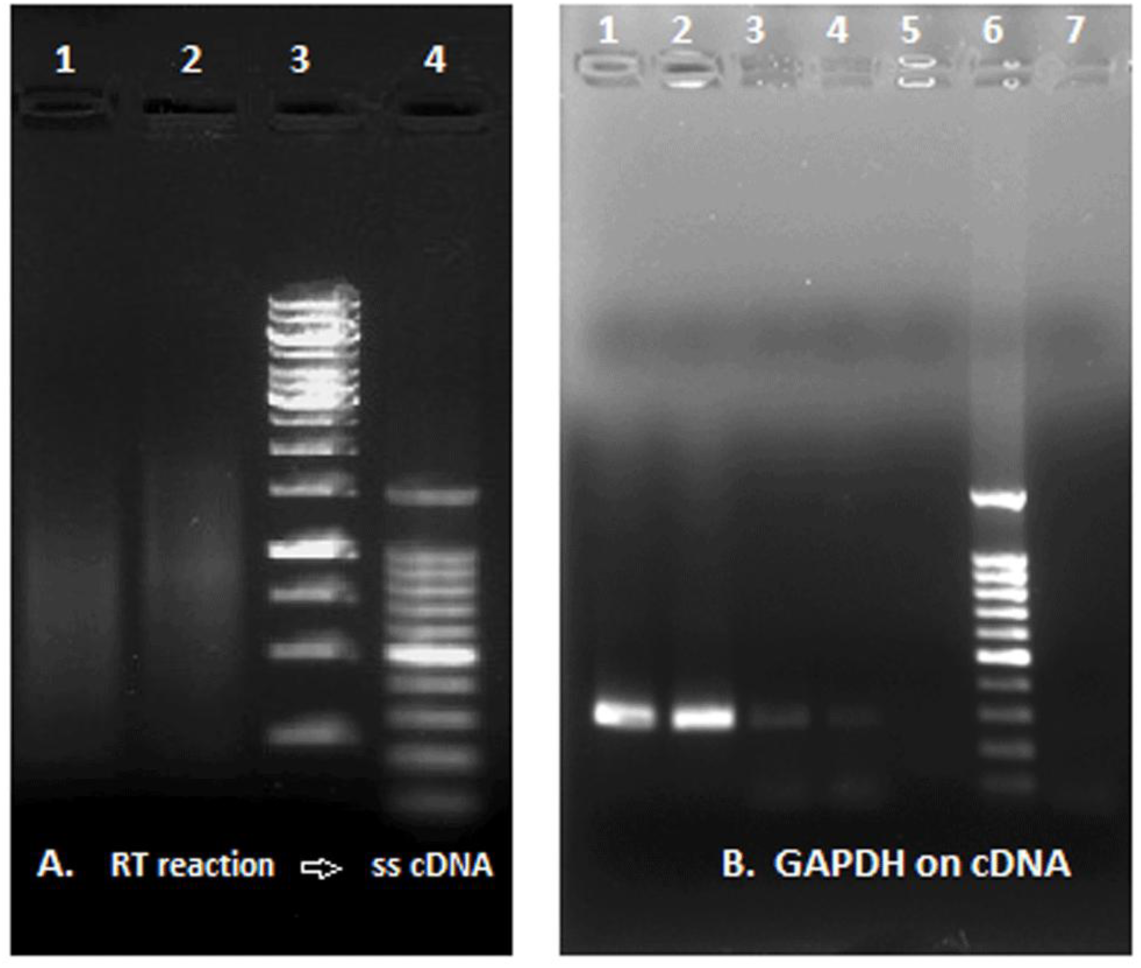
Reverse Transcriptase (RT) activity of the human Retro-Giant viruses. (A) RT reaction and synthesis of ss c-DNA: Lane 1, reaction with a commercial RT enzyme; Lane 2, reaction with viral pellet (reaction without RT enzyme); Lane 3, Gene Ruler 1Kb DNA ladder; Lane 4, 100bp DNA ladder. (B) GAPDH amplification from ss-cDNA template: Lanes 1 and 2, reaction with commercial RT enzyme; Lanes 3 and 4, reaction with the lysed viral pellet; Lane 5, negative control; Lane 6, DNA ladder; Lane 7, additional negative control.

**Figure 5.**
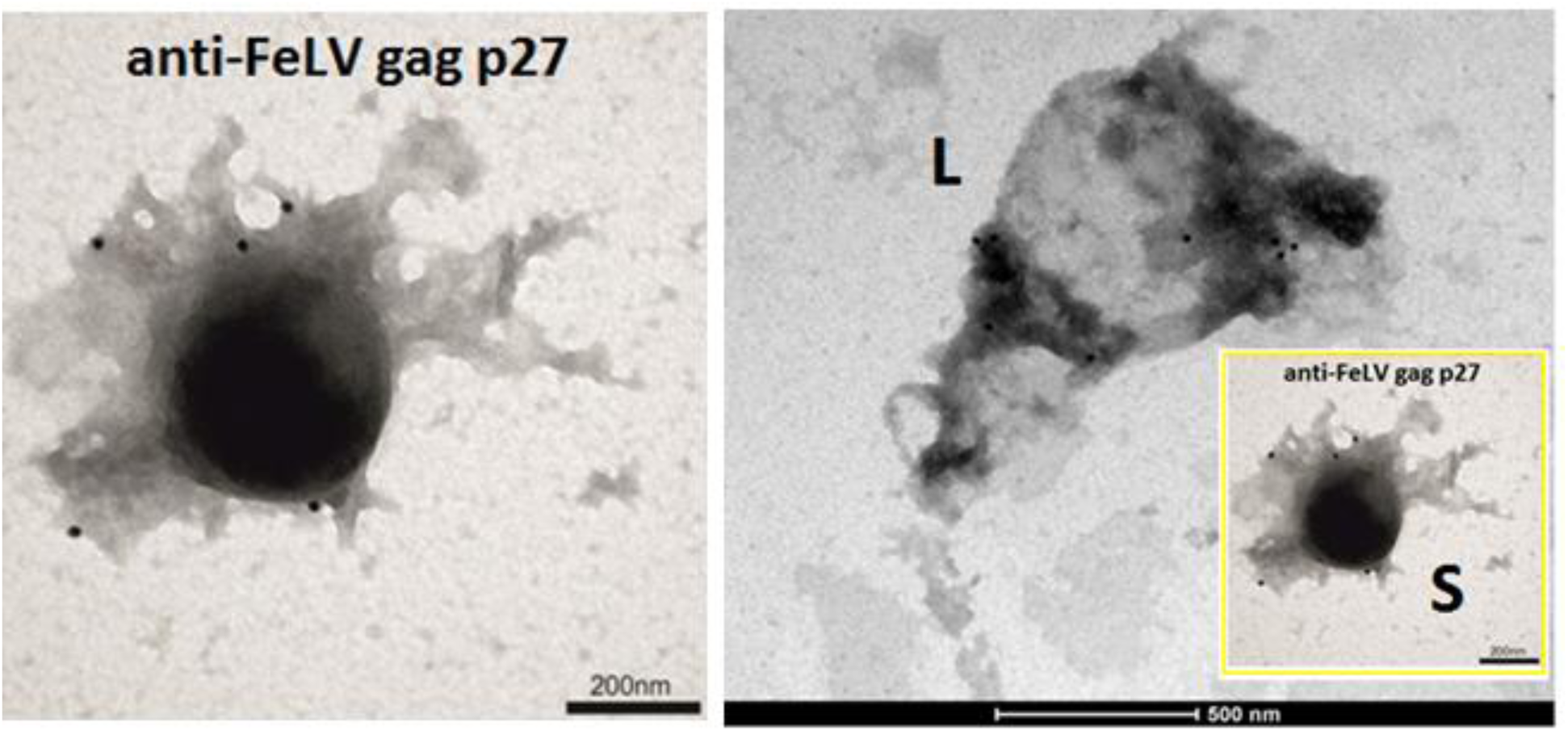
The large ∼1 μm particle and the smaller satellite ∼400-450 nm giant viral particles share the same retroviral antigenicity, being both specifically labelled by anti-FeLV gag p27 MoAb.

### Whole genome shotgun sequences

The assembly of the 75bp paired-end reads, derived from the shotgun sequencing of this microbial agent viral entity triggered to the production of a partial genome (scaffold) of 1.48MB. The algorithm used was Blast2GO that disclosed a mega-genome of 1.48 MB (Total length 1481106). This is the size length that is typical of mega-genomes also reported for environmental giant viruses ^17^.

In the initial step of gene prediction, we roughly discovered 1116 putative ORFs, reduced to ∼1023 after the application of BLAST2GO software for gene similarity finding. The products of such predicted genes have affinity with various genes found in different domains of life. In particular, we discovered many features related to Archaea, (thermosome, anaerobic metabolism, operons, histone-like proteins, Archaea topoisomerase and. translation initiation factor), showing a predominance of this domain within this microbial entity. The annotated Archaea features by bioinformatics are consistent with the morphology of the large particle that looks like an ancestral unicellular organism. Nevertheless, the assembled genome cannot be completely associated to the Archaea domain as 16S rRNA has not been identified.

Substantially, BLAST2GO algorithm reconstructed an ancestral self-governing microbial cell-like entity with genes in common to the three domains of life, with a complex system of genes that regulate cell proliferation and epigenetics. The concept of microbial cell-like entity, with genes in common to the three domains of life, also applies to environmental giant Mimiviruses, entities, for some aspects, more closely to a cell than to a virus ^18^.

Simultaneously, we detected interesting retroviral genes, including those codifying the *gag-pol* complex that explained the retro-transcriptase activity in vitro, and the oncogenic *gag-akt* retroviral kinase. The latter is highly related to FELV (E-value 2.00E-33; >95.04% identity), confirming the retroviral antigenicity documented at the EM immunogold.

Thus, the fascinating chimeric nature of this transforming microbial entity not only relies on a retroviral nature, but its large genome discloses a real cancer factory, with a complex system of oncogenes.

The mega-genome displays sequences with a significant homology with the human *c-myc* gene, *v-myb* oncogene and a set of serine-threonine protein kinases, very similar to the eukaryotic ones. In addition, Cyclin dependent kinases (CDKs) and cells division proteins have been predicted to be part of this microbial giant agent.

Most of these features representing the complex chimeric nature of this microbial system and supporting its capability to promote cancerogenic processes are highlighted in Figure 4, where the whole BLAST2GO gene annotation is schematized.

The whole genome sequence is reports in the *Annotation GO file* https://drive.google.com/open?id=1ZjCc4aEsEVvJO-98z_hPpaJ6ZwKG2zt2; FastQ and FastQC in the following link https://mega.nz/#F!7dID0aSa!8bA-4qVdPeiY0tsSbd8G7g.

**Figure 4.**
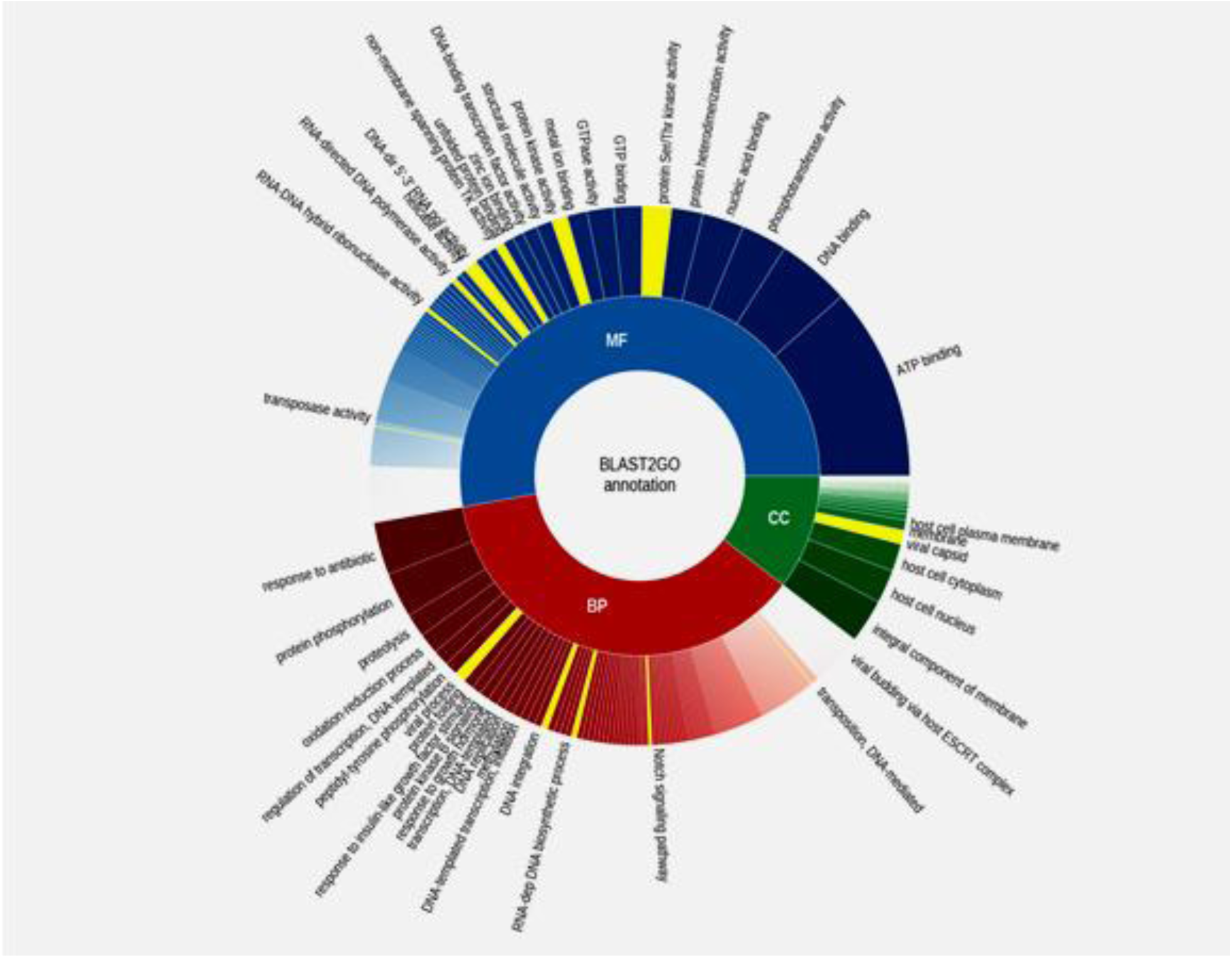
BLAST2GO algorithm reconstructed an ancestral self-governing microbial entity with genes in common to the three domains of life, with a complex system of genes that regulate cell proliferation and epigenetics.

## DISCUSSION

Robert Gallo reported the first human retrovirus HLTV in 1980 ^19^. What we report here is the discovery of the first Mimivirus-sized human giant virus with oncogenes and a transforming retroviral kinase.

In our previous work, conducted initially on human tissues, EM depicted previously unreported ∼400-450 nm gigantic particles associated with large aggregates, resembling viroplasms, recognized by anti-FeLVp27 gag Ab ^1^. The particle diameters were more than four times the 100 nm size expected in retroviruses. These large particles and associated structures discovered in human cells appeared to morphologically parallel previously reported amoeba Mimiviruses ^6^. Gram positive blue granules that disclosed the existence of giant viruses in the amoeba, similarly detected this newly giant agent in human cells. Proteomic analyses suggested the presence of histone H4 variants common to environmental giant DNA viruses. However, the striking difference was the unique mammalian retroviral nature of the human giant particles. Distinct from the amoebas’ Mimiviruses, the viral particles were immuno-labelled with anti FeLV p27gag moAb.

How to prove if we were really facing giant microbial entities with a retroviral nature? Working on human tissues was confusing and the distinction between the virus and the cells became blurred. Therefore, in order to distinguish the giant agent from human cells, we attempted to isolate the viral particles through a sucrose gradient.

A white ring was detected in the 25% sucrose fraction - the same sedimentation fraction that is also typical of giant Mimiviruses isolated from amoeba ^6^. Electron Microscopy confirmed the presence of giant viral particles. The harvest from human cancer cells of these giant microbial entities on sucrose became a routine test, but it did not require amoeba co-culture. Our particles were not *rumour viruses* inside tissues anymore, but microbes that can be harvested in few hours and stored as a pure microbial aliquot. Every time, the particles were tested for their RT activity and they had reverse transcriptase activity.

An in depth morphological analyses revealed that the predominant 400-450 nm giant viral particles, surround a much larger giant particle. These human biological entities appear as a microbial system composed by a large particle of ∼1-1.5 micron, strongly recalling ancestral unicellular organisms, and smaller satellite associated viral particles, Mimiviruses-sized. The large particle can reach up to 2.0 μm.

The shotgun sequencing triggered to the production of a partial genome scaffold of 1.48 MB, lenght that it is typical for giant microbes ^17^. Blast2GO discovered many Archaea’s features, showing a predominance of this domain within this new microbial entity. Nevertheless, the assembled genome cannot be completely associated to the Archaea domain, as 16S rRNA has not been identified.

Simultaneously, we detected interesting retroviral genes, including those codifying the transforming retroviral kinase *gag*-*akt fusion protein*, related to feline retrovirus lineage ^20-21^. This confirmed the specific immuno-labelling of the giant viral particles with a screen of anti-FeLV Mo-Abs and the retrotranscriptase enzymatic activity of the giant viral particles.

For their giant dimensions, analogies with environmental Mimiviruses and the retroviral nature that complete their chimeric essence, we tentatively named these biological entities Retro-Giant Viruses (RGV). However, distinct from Mimiviruses infecting amoeba, that display a different genomes and tropism ^22-23^, RGV have oncogenes and a transforming retroviral kinase and do not require amoeba co-culture, since amoeba is not their natural host.

The fascinating nature of the RGV not only relies on their retroviral nature, but their large genome discloses a real cancer factory with a complex system of oncogenes with the archaeal ancestry. We found many evidences of “archaeal oncogenes”. The RGV display sequences with a significant homology with the human *c-myc* gene product, which in turn shares common epitopes with an archaebacterial protein from *Halobacterium halobium* ^*24*^. RGV regions homologous to the *v-myb* oncogene are also found in both halophilic and methanophilic archaebacteria ^25^. The set of serine-threonine protein kinases, listed in our functional annotations, are very similar to the eukaryotic ones, but simultaneously homologous to those identified in some archaebacteria including Methanococcus vannielii, M. voltae and M. Thermolitotrophicus ^26-28^.The predicted RGV helicase is an archaebacterial protein involved in the tumour-suppressor pathways ^29^. In addition, cyclin dependent kinases (CDKs) and cells division proteins, playing an essential role in the common archeal cell cycle, have been predicted to be part of this of RGV genome ^30^.

The Retro-Giant viruses represent unique microbial cell-like entities with genes in common to the three domain of life where a retrovirus is intrinsically present. In the transition from the RNA to DNA world, three different viruses —the RNA virus, the DNA virus, and a retrovirus—co-infected a cell. This scenario describes the essence of the large particle that resembles ancestral unicellular organisms ^31^. It will remain to define the role of the large particle and the satellite giant viruses in disseminating oncogenes.

Blast2GO algorithm provided a good example of realistic algorithms that closely reflected the real life scenario, linking the bioinformatics scores with reproducible biological evidences and tumour formation in mice ^32^.

Fulfilling Koch’s postulates, we demonstrated how constant was the isolations from human cancer cells of these giant viral particles and how to use them to transform normal cells. The subsequent re-extraction of the giant particles from transformed infected NIH 3T3 cells and tumour explants from infected mice, confirmed the transmissibility of this human agent.

The morphology, biology and genetic features allocate this mammalian giant microbe halfway in between a classic oncogenic virus and an infectious cancer cell. The complexity of its nature makes the agent a sort of small autonomous infectious cell, a simplified version of its eukaryotic counterpart, specialized in carcinogenesis. RGV constantly had the ability to induce tumours formation in mice and this was in virtue of their oncogenes and transforming gag-akt retroviral kinase. A filtered supernatant did not transform and this ruled out the presence of filterable retroviruses.

## Conclusion

Science grew with the assumption that viruses were small and filterable, but the discovery of Mimiviruses infecting amoeba modified this perception, revealing microbial agents of exceptional size and genome complexity. Mimiviruses highlighted how the classical virus isolation procedures, that include filtration through a 0.45 μm filter, may prevent the recovery of large microbes. In virtue of this paradigm change, scientists were provided with the right tools to interpret and discover a variety of microbes that resist current classifications ^33^. Despite being substantially different from Mimiviruses infecting protists, the newly identified oncogenic giant microbe that infect humans and animals is the corollary of this revolution. The common filtration techniques missed an entire shuttling system of oncogenes enclosed in an infectious unicellular life form left undisturbed for centuries.

## Graphic Summary

https://drive.google.com/open?id=1rcYI8Ex3rDyBN-lVukBYaxYNm-xJ6b9n

## Data availability

All slides, entire electron microscopy collection and stored microbial aliquots are available to be examined; please contact the corresponding author.

## Acknowledgements

The anti-FeLV-related mo-Abs were kindly provided as a gift by Dr Chris Grant of Custom Monoclonals International (West Sacramento, CA 95691, USA).

